# Elucidating the JNK Signaling Pathway in Neonatal Muscle Growth and Neuromuscular Contractures

**DOI:** 10.64898/2026.06.30.735638

**Authors:** Kasey Shao, Madelyn Shoates, Daniela Barrios, Sophia Conte, Albaraa Tarabishi, Geethika Velaga, Kritton Shay-Winkler, Qingnian Goh, Roger Cornwall

## Abstract

Neuromuscular contractures arising from neonatal brachial plexus injuries (NBPI) are highly disabling and currently incurable. We previously showed that contractures involve impaired longitudinal growth of denervated muscles, a defect mediated through myostatin (MSTN) signaling, a potent negative regulator of muscle size. However, MSTN-mediated contractures occur independent of canonical signaling pathways, including SMAD 2/3 and AKT/mTOR. Through a mouse model of NBPI, our present study extended these findings by revealing pharmacologic inhibition of JNK signaling, a noncanonical pathway downstream of MSTN, partially rescues contractures without restoring muscle length. Rather, JNK activation upregulates myofiber expression of the target gene *Lmna*, which encodes the nuclear envelope proteins Lamin A and Lamin C that are vital for nuclear stability, resulting in pervasive myonuclear displacement. These results suggest that other factors contribute to contracture pathology beyond deficits in longitudinal muscle growth. Further, while JNK inhibition does not restore length of denervated muscles, it impedes size and mass of normally innervated neonatal muscles, suggesting a requirement of JNK signaling for neonatal muscle growth. Our collective findings thereby establish new mechanistic insights into the molecular basis of aberrant muscle growth and neuromuscular contracture formation, potentially leading to novel targets for muscle restorative strategies and medical contracture prevention.

## Introduction

Neonatal Brachial Plexus Injury (NBPI) is the most common cause of upper extremity paralysis in children, occurring in 1 to 3 per 1,000 live births.^1^ Although the paralysis can recover, this initial nerve injury results in the secondary formation of disabling muscle contractures, which cannot be prevented or cured with existing therapies.^2,3^ Instead, contractures severely reduce joint range of motion and limit functional use of the affected limb, ultimately leading to progressive skeletal deformities.^4–7^ Using a mouse model of NBPI, we first showed that contractures involve impaired longitudinal growth of denervated muscles, as characterized by sarcomere overstretch.^8–11^ We and others have since observed this finding clinically in children and infants,^12–14^ as well as replicated it in similar rat models.^15–17^ We next discovered this deficit in muscle length is modulated by increased proteasome-mediated protein degradation,^18,19^ as systemic administration of proteasome inhibitors following NBPI can rescue longitudinal muscle growth and prevent contractures.^18–20^ However, the off-target toxicity of proteasome inhibition calls for investigation of muscle-specific therapies.^18,20^ Using a soluble ligand trap, we found that inhibition of myostatin (MSTN), a muscle-specific inhibitor of muscle growth,^21,22^ can reduce proteasome activity, restore longitudinal muscle growth, and prevent contractures after NBPI.^23^ However, the contracture rescue with pharmacologic MSTN inhibition is incomplete and not associated with alterations in canonical MSTN signaling pathways (SMAD 2/3, Akt/mTOR).^23^ To develop novel targets for contracture therapy, we must continue to interrogate the mechanistic role of MSTN signaling in contracture pathogenesis by pivoting to its non-canonical intracellular pathways, including Ras-MEK1-ERK1/2 and TAK1-JNK-c-Jun.^24^

Ligand binding of MSTN to its Type 2 receptor, ACVR2B, initiates Extracellular Receptor Kinase (ERK) signaling through activation of RAF kinases, and subsequently mitogen-activated protein kinase kinase 1 (MEK1), which in turn phosphorylates and activates ERK1/2.^25,26^ The ERK pathway has pleiotropic roles in muscle health and function, including myogenic differentiation, muscle mass and fiber-type maintenance, as well as disease modification in muscular dystrophies.^27–30^ In comparison, the c-Jun N-Terminal Kinase (JNK) pathway attenuates skeletal muscle growth and is downregulated during myogenesis.^31^ MSTN binding to ACVR2B initiates JNK signaling through activation of TGF-B-activated kinase-1 (TAK1), and subsequently the mitogen-activated protein kinase (MAPK) family member MKK4, which then activates JNK.^31,32^ This signaling cascade elicits various growth inhibitory pathways, including the activation of the caspase-8/caspase-3 signaling cascade that induces apoptosis,^33^ and phosphorylation of the c-Jun protein, which upregulates *DNMT1* expression and promotes genome-wide methylation.^34^ Additionally, phosphorylated c-Jun heterodimerizes with the c-Fos protein to form the transcriptional factor, Activating Protein 1 (AP-1), which drives transcription of an array of target genes.^35^ Therefore, in this study, we sought to first elucidate the role(s) of ERK and JNK signaling in contracture formation. Additionally, given the breadth of potential effectors regulated through these pathways, we sought to further identify the downstream target(s) modulating contractures. Our findings here show that NBPI-induced contractures involve upregulation of the JNK signaling pathway and its downstream target *Lmna*. Furthermore, we identified that contractures involve factors beyond impaired longitudinal muscle growth, including myonuclear dysmorphia and displacement. These collective results thereby advance our mechanistic understanding of the pathophysiology of neuromuscular contractures.

## Results

### JNK signaling modulates contracture formation

To elucidate the role of non-canonical MSTN signaling pathways in contracture formation, we began by characterizing ERK and JNK activity in neonatally denervated muscles (**Fig 1A, 1E**). Throughout four weeks of NBPI, levels of ERK1/2 phosphorylation and ERK1/2 signaling (defined as ERK1/2 activation normalized to ERK1/2 translation) remain unelevated, despite an increase in ERK1 phosphorylation and in total ERK and ERK1 protein translation (**Fig B-D, SFig 1A-F**). Conversely, JNK1/2 phosphorylation and JNK1/2 signaling (JNK1/2 activation normalized to JNK1/2 translation), as well as levels of phosphorylated JNK1 and JNK2 isoforms, and JNK1 and JNK2 signaling, are elevated in denervated muscles at two weeks post-NBPI (**Fig 1F-H, SFig 1G-L**), a timepoint where elbow contractures are beginning to form in mice.^11^ We next assessed a potential role for this upregulation in JNK activity in modulating contractures by incorporating the pharmacological inhibitor, SP600125 (selective inhibitor of JNK1/2),^36,37^ with our surgical NBPI mouse model (**Fig 2A**). Treatment with SP600125 for two weeks after NBPI attenuates the increase in JNK1/2 phosphorylation in denervated muscles, restoring JNK1/2 signaling to levels comparable to non-operated muscles (**Fig 1E-H**). Of interest, this inhibition in overall JNK activity corresponds specifically to a reduction in JNK1 (but not JNK2) phosphorylation and signaling (**SFig 1G-L)**. Critically, at four weeks post-NBPI, pharmacologic JNK inhibition partially restores elbow joint motion and rescues elbow contractures (**Fig 2B-D**). These collective results thereby indicate that contractures are at least partially modulated through the JNK signaling pathway, and preferentially through JNK1 signaling. Moreover, they allow us to eliminate the ERK pathway and direct our focus to dissecting the intracellular mechanisms and downstream effectors of the JNK cascade in contracture pathogenesis.

**Figure 1:**
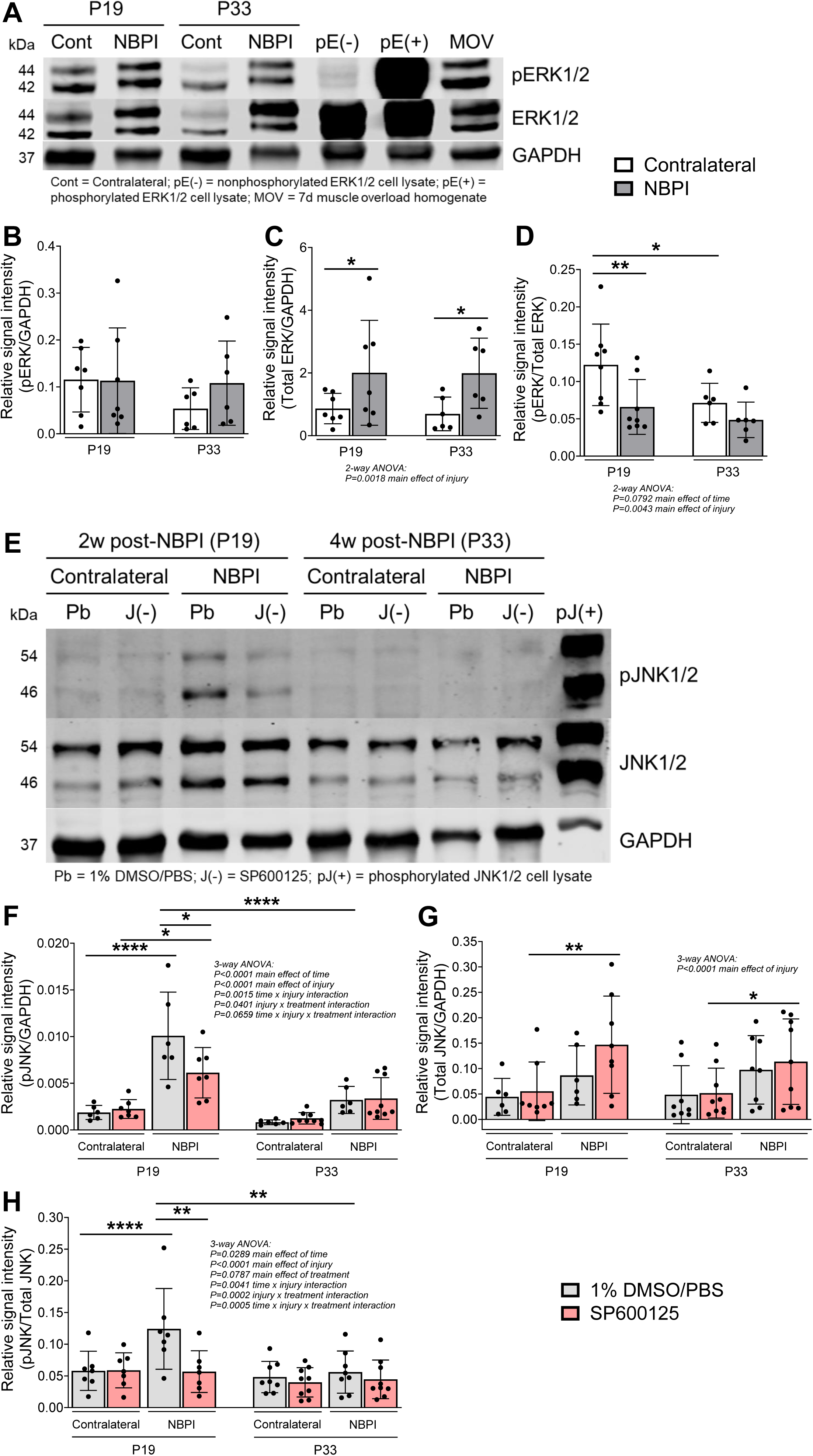
Neonatal muscle denervation activates JNK signaling. (**A**) Representative western blots and quantitative analysis of (**B**) phosphorylated ERK1/2 (pERK), (**C**) total ERK1/2, and (**D**) the western signal for pERK1/2 normalized to total protein levels indicated that while neonatal denervation increases ERK translation in biceps brachii muscles, both ERK activation and signaling are not elevated throughout four weeks of NBPI. Data are presented as mean ± SD, n = 6-8 independent mice per group. In contrast, (**E**) representative blots and quantitation of (**F**) phosphorylated JNK1/2 (pJNK), (**G**) total JNK1/2, and (**H**) the normalized pJNK1/2 western signal revealed an upregulation of both JNK activation and signaling in denervated muscles, specifically at two weeks post-NBPI. Furthermore, pharmacologic JNK inhibition with SP600125 (**F**) attenuates the increase in JNK phosphorylation, and (**H**) restores JNK signaling to baseline levels in neonatally denervated muscles. Molecular weights: ERK1 (44 kDa), ERK2 (42 kDa), JNK1 (46 kDa), JNK2 (54 kDa). Data are presented as mean ± SD, n = 6-9 independent mice per group. Statistical analyses: (**B**), (**C**), (**D**) two-way ANOVA for NBPI surgery (repeated measures between forelimbs) across time with a Bonferroni correction for multiple comparisons, (**F**), (**G**), (**H**) three-way analysis of variance (ANOVA) for pharmacologic treatment and NBPI surgery (repeated measures between forelimbs) across time with a Bonferroni correction for multiple comparisons. **\***p < 0.05, **\*\***p < 0.01, **\*\*\*\***p<0.0001.

**Figure 2:**
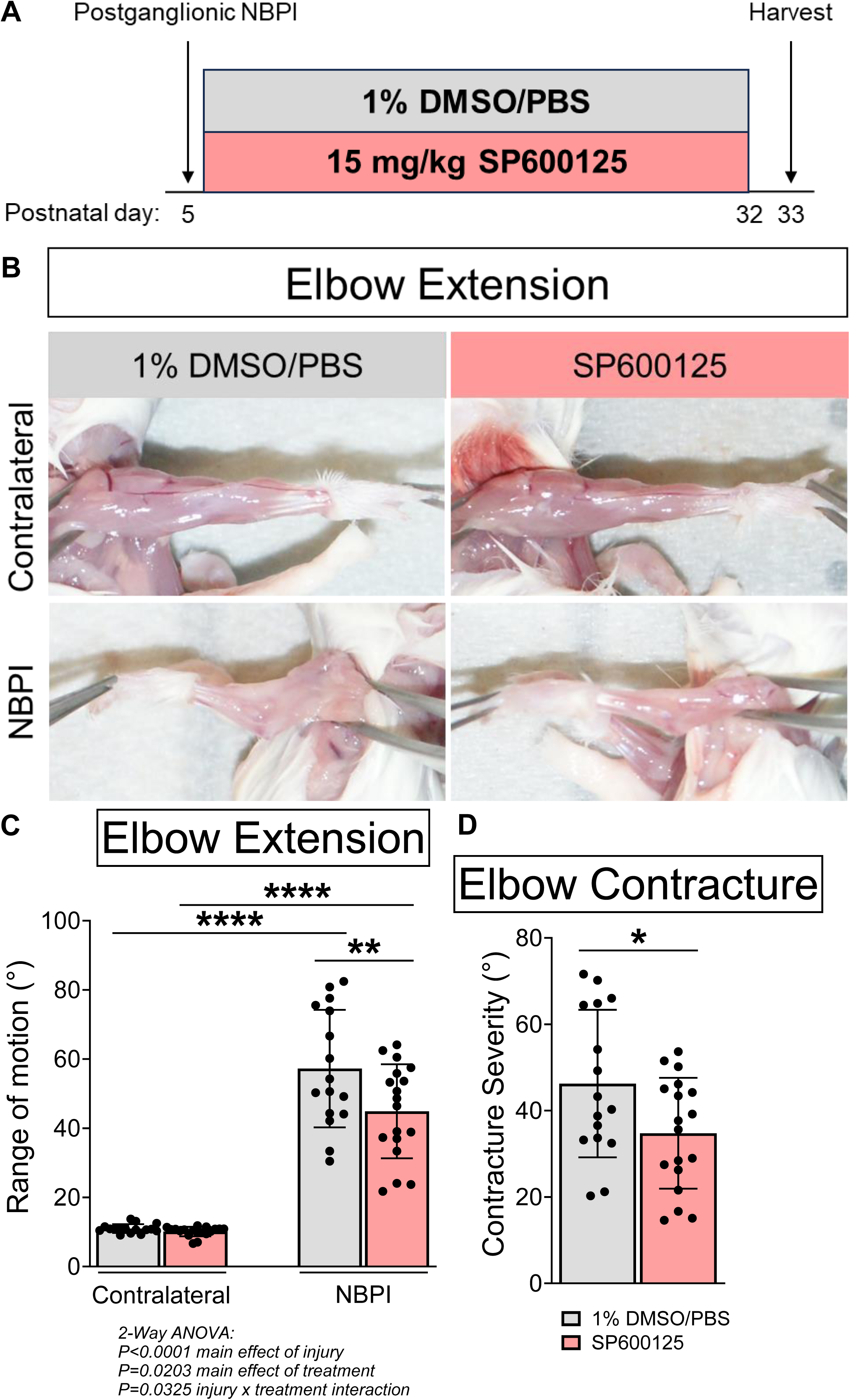
Pharmacologic JNK inhibition reduces NBPI-induced contractures. (**A**) Schematic depicting 1% DMSO/PBS (vehicle) or 15 mg/kg SP600125 treatment to pharmacologically inhibit JNK signaling in mice during neonatal muscle growth (P5 – P32), and assessment of joint motion at P33 (four weeks post-NBPI). (**B**) Representative images of denervated (NBPI) and non-operated (contralateral) forelimbs, and quantitative analysis of both (**C**) passive elbow extension and (**D**) elbow flexion contracture severity revealed that pharmacologic JNK inhibition prevents the formation of elbow contractures. In (**D**), elbow flexion contracture severity is calculated as the difference in passive elbow extension between the NBPI and contralateral sides. Data are presented as mean ± SD, n = 16-18 independent mice per group. Statistical analyses: (**C**) two-way ANOVA for pharmacologic treatment and NBPI surgery (repeated measures between forelimbs) with a Bonferroni correction for multiple comparisons, (**D**) unpaired two-tailed Student’s t-tests between treatment groups. *p<0.05, **p<0.01, ***p<0.001, ****p<0.0001.

### JNK signaling is required for neonatal muscle growth

We next extended our investigation of this pathway to various dimensions of muscle growth, given the known role of longitudinal muscle growth in contracture development. At four weeks post-NBPI, pharmacologic JNK inhibition fails to protect denervated muscles against sarcomere overstretch, indicating the rescue in contractures occurs in the absence of functional improvements in longitudinal muscle growth (**Fig 3A-C**). Critically, SP600125 treatment creates a tighter correlation between contracture severity and sarcomere overstretch, indicating the removal/improvement of other elements of contracture pathology that do not pertain to muscle length (**Fig 3D-E**). Similar to length, pharmacologic JNK inhibition does not restore cross-sectional area, volumetric growth, or overall mass of denervated muscles (**Fig 4A-D**). Rather, it reduces muscle volume and mass only in normally innervated muscles (**Fig 4C-D**). These deficits in neonatal muscle growth were accompanied by reductions in whole body weight gains, but not humerus lengths (**SFig 4A-B**); thus SP600125 induces muscular but not skeletal growth defects. Taken together, our findings suggest that while JNK signaling is an essential regulator of muscle growth during the neonatal period, it modulates contractures independent of alterations to muscle growth.

**Figure 3:**
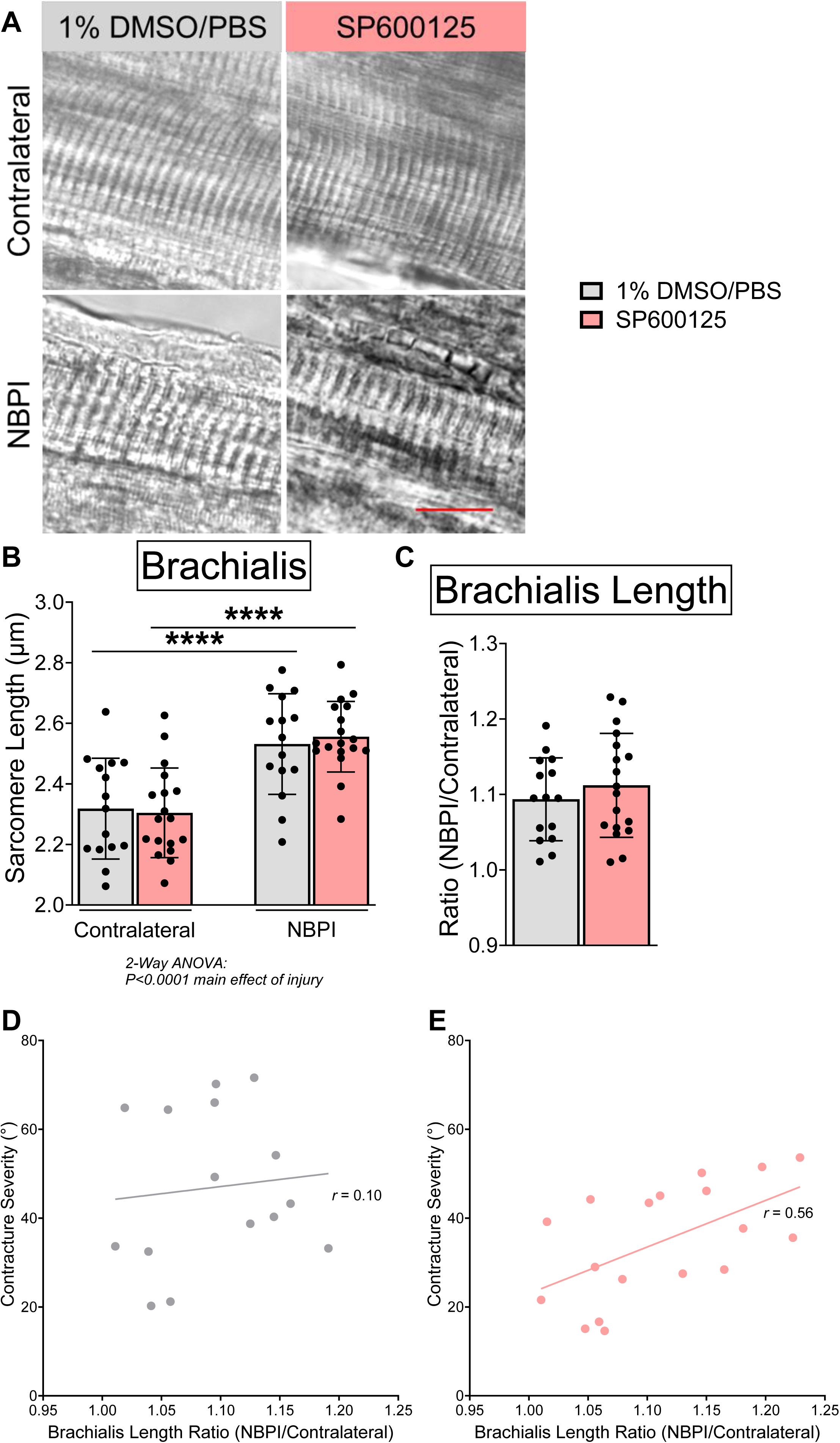
Pharmacologic JNK inhibition does not protect against sarcomere overstretch following NBPI. (**A**) Representative DIC images of sarcomeres, and quantitative analysis of (**B**) absolute sarcomere length and (**C**) sarcomere length ratio in brachialis muscles showed that SP600125 treatment does not reduce sarcomere overstretch after NBPI, indicating the rescue in elbow contractures with pharmacologic JNK inhibition occurs independent of improvements to functional muscle length. In (**C**), sarcomere length ratio is calculated by normalizing sarcomere length in the denervated forelimb (NBPI) to its contralateral side, as an assay for functional muscle length. Within mice, pairing elbow contracture severity with the corresponding sarcomere length ratio revealed a (**D**) scattered cluster with vehicle treatment, and a (**E**) tighter cluster with SP600125 treatment, indicating JNK inhibition removed factors not relevant to muscle length. Data are presented as mean ± SD, n = 15-18 independent mice per group. Statistical analyses: (**B**) two-way ANOVA for pharmacologic treatment and NBPI surgery (repeated measures between forelimbs) with a Bonferroni correction for multiple comparisons, (**C**) unpaired two-tailed Student’s t-tests between treatment groups. (**D**), (**E**) Pearson’s correlational analysis between elbow contracture severity and sarcomere length ratio within mice. ****p<0.0001. Scale bar (**A**): 10 µm.

**Figure 4:**
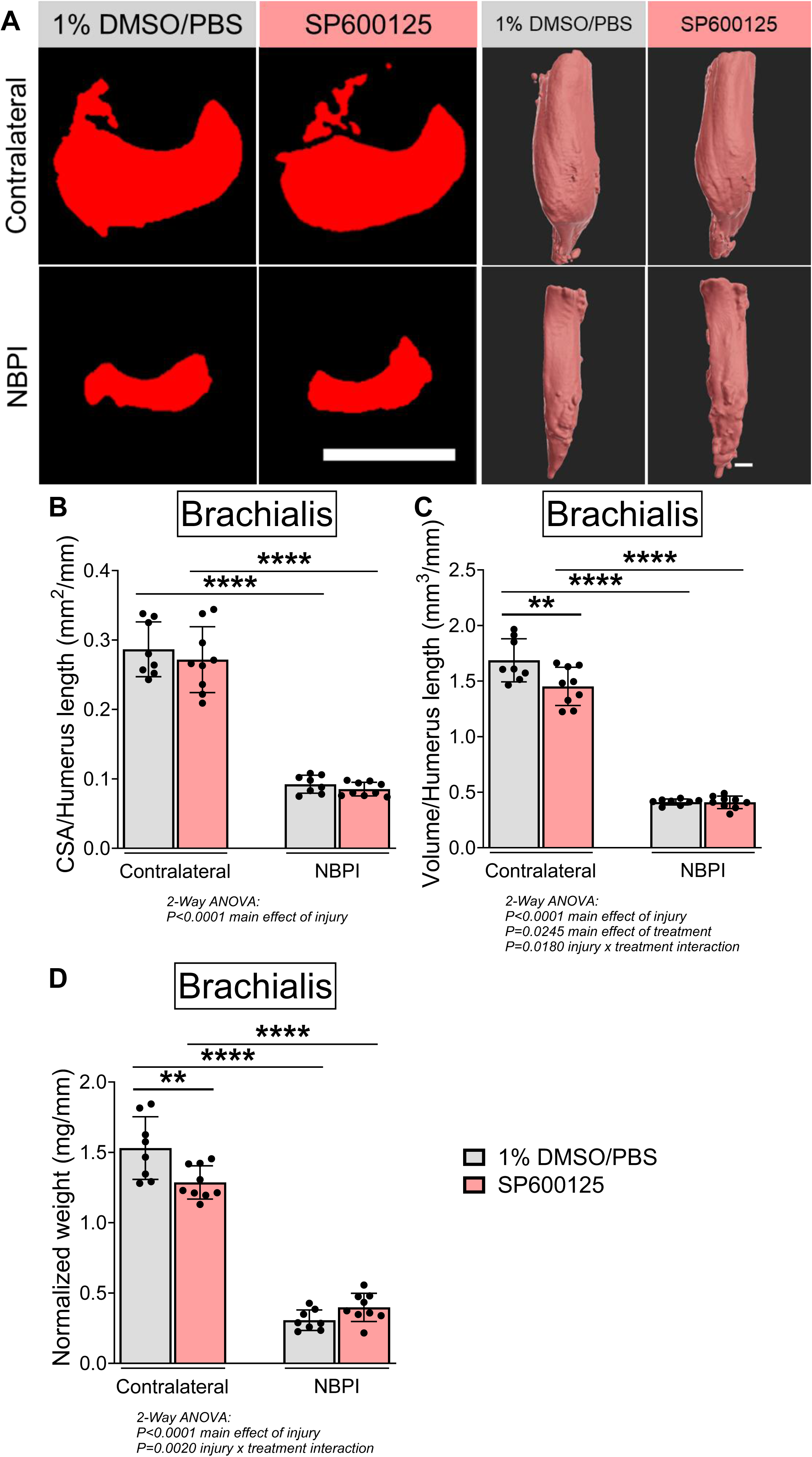
Pharmacologic JNK inhibition blunts neonatal skeletal muscle growth. (**A**) Representative micro-CT images of brachialis muscles in transverse and 3-dimensional views, and quantitative analysis of whole muscle (**B**) CSA and (**C**) volume revealed smaller muscles in normally innervated (contralateral) forelimbs with SP600125 treatment. (**D**) Neonatal JNK inhibition further reduces the weight of non-denervated brachialis muscles. Data are presented as mean ± SD, n = 8-9 independent mice per group. Statistical analyses: (**B**), (**C**), (**D**) two-way ANOVA for pharmacologic treatment and NBPI surgery (repeated measures between forelimbs) with a Bonferroni correction for multiple comparisons. **p<0.01, ****p<0.0001. Scale bars (**A**): 1000 µm.

### Lmna is a potential downstream effector of JNK-mediated contractures

We next sought to identify downstream pathways by which JNK-dependent contractures occur. We began by examining known downstream targets of JNK signaling, which include increased apoptosis driven by elevated Caspase-3 levels, increased DNA methylation driven by upregulated *Dnmt1* expression, and altered expression of gene targets of AP-1. At 4 weeks post-NBPI, protein levels of Caspase-3 and transcript levels of *Dnmt1* are upregulated in denervated muscles, but are not reduced with pharmacologic JNK inhibition (**Fig 5A-B**). Moreover, neonatal denervation alone does not upregulate the active (cleaved) form of Caspase-3, which serves as the specific biomarker for dying or dead cells. Thus apoptosis and DNA methylation are unlikely pathways by which JNK signaling mediates contractures. We next examined six downstream target genes with putative AP-1 binding sites:^38^ *Atl1*, *Fam126a*, *Fgd4*, *Flnc*, *Kif1a*, and *Lmna*, all of which have known human muscle phenotypes by gene ontology.^39,40^ Neither neonatal muscle denervation nor SP600125 alters transcript levels of *Atl1*, *Fam126a*, *Fgd4*, *Flnc*, and *Kif1a* (**Fig 5C-G**). In contrast, NBPI upregulates *Lmna* mRNA expression and Lamin A/C protein levels, both of which are attenuated by pharmacologic JNK inhibition (**Fig 5H-I**). These findings point to *Lmna* as a promising target of JNK in contracture modulation.

**Figure 5:**
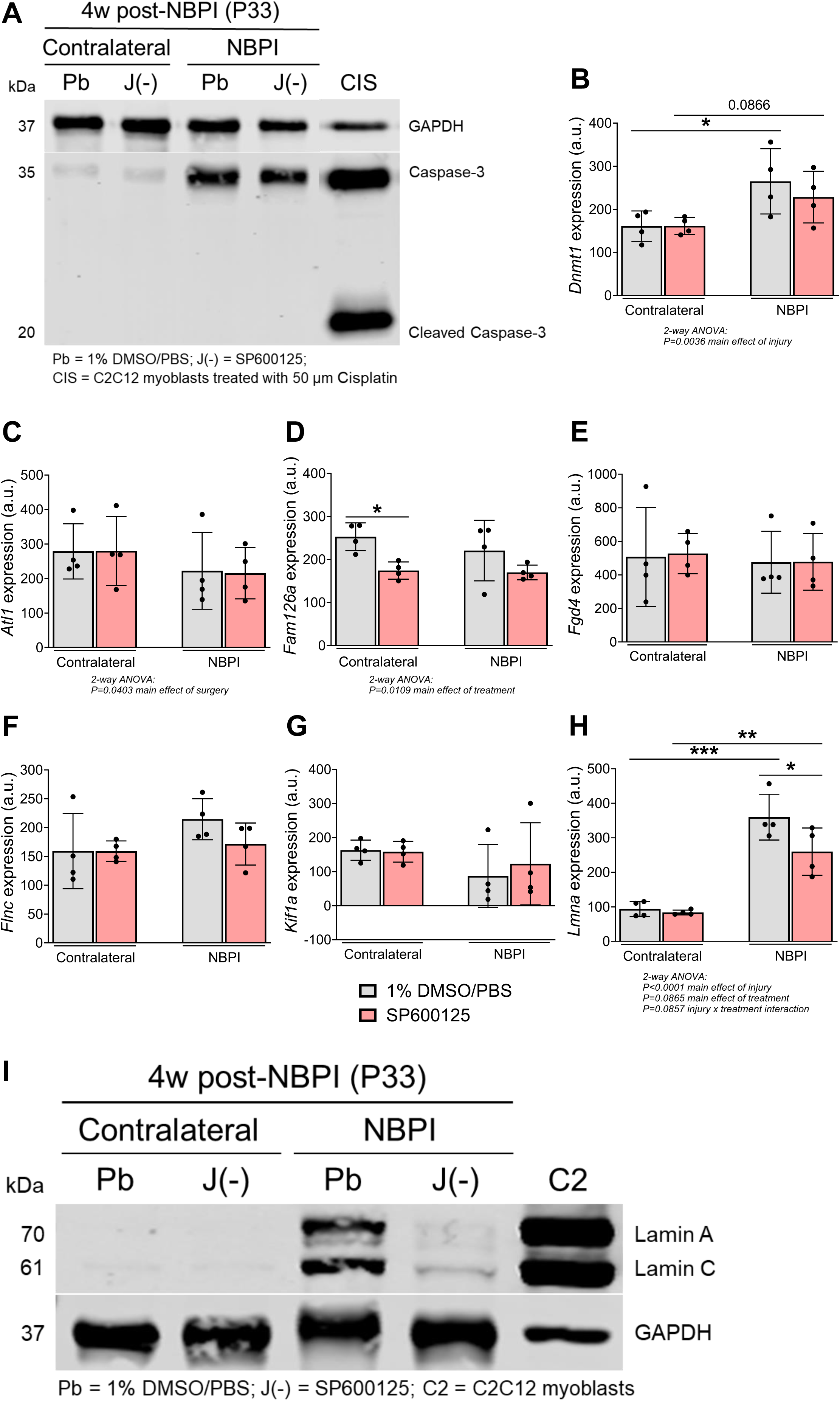
Pharmacologic JNK inhibition attenuates *Lmna* expression following NBPI. (**A**) Protein levels of Caspase-3, and transcript levels of (**B**) *Dnmt1*, (**C**) *Atl1*, (**D**) *Fam126a*, (**E**) *Fgd4*, (**F**) *Flnc*, and (**G**) *Kif1a* in biceps muscles are not altered with denervation and/or SP600125 treatment. In contrast, JNK inhibition attenuates the elevated levels of (**H**) *Lmna* mRNA transcription and (**I**) Lamin A/C protein translation in denervated biceps muscles after NBPI. Data are presented as mean ± SD, n = 4 independent mice per group. Statistical analyses: (**B**), (**C**), (**D**), (**E**), (**F**), (**G**), (**H**) two-way ANOVA for pharmacologic treatment and NBPI surgery (repeated measures between forelimbs) with a Bonferroni correction for multiple comparisons. *p<0.05, **p<0.01, ***p<0.001.

To gain insights on the contributions of *Lmna* to contracture pathophysiology, we began by further characterizing its expression during NBPI. Immunostaining of normally innervated muscles revealed minimal Lamin A/C protein levels, primarily in nuclei residing outside of myofibers, as well as in myonuclei residing within the periphery of myofibers (**Fig 6A-B**). Conversely, in denervated muscles after NBPI, substantial levels of Lamin A/C protein can be detected in spherical, centrally located myonuclei stacked longitudinally within the myofiber (**Fig 6A-B**). To complement our findings of elevated Lamin A/C levels during conditions of aberrant muscle growth, we utilized C2C12 myoblasts to characterize temporal *Lmna* expression during in vitro myogenesis. Western blot detected an upregulation of Lamin A/C protein level during myoblast proliferation, which declines throughout muscle differentiation (**SFig 6A**). To further explore *Lmna* function, we overexpressed the Lamin A and Lamin C isoforms of *LMNA* in separate cultures of myoblasts at 24h post-differentiation (**SFig 6B**). After three days (72h) of differentiation, dysmorphic changes were observed in Lamin A and Lamin C overexpressing cultures, resulting in wider and rounder multinucleated muscle cells than cultures expressing an empty vector (Empty) (**Fig 6C-G**). These results thereby posit an intriguing association between contractures, *Lmna* overexpression, and altered myonuclear morphology and positioning *in vivo*, as well as a role for Lamin A/C in guiding myocyte morphology *in vitro*.

**Figure 6:**
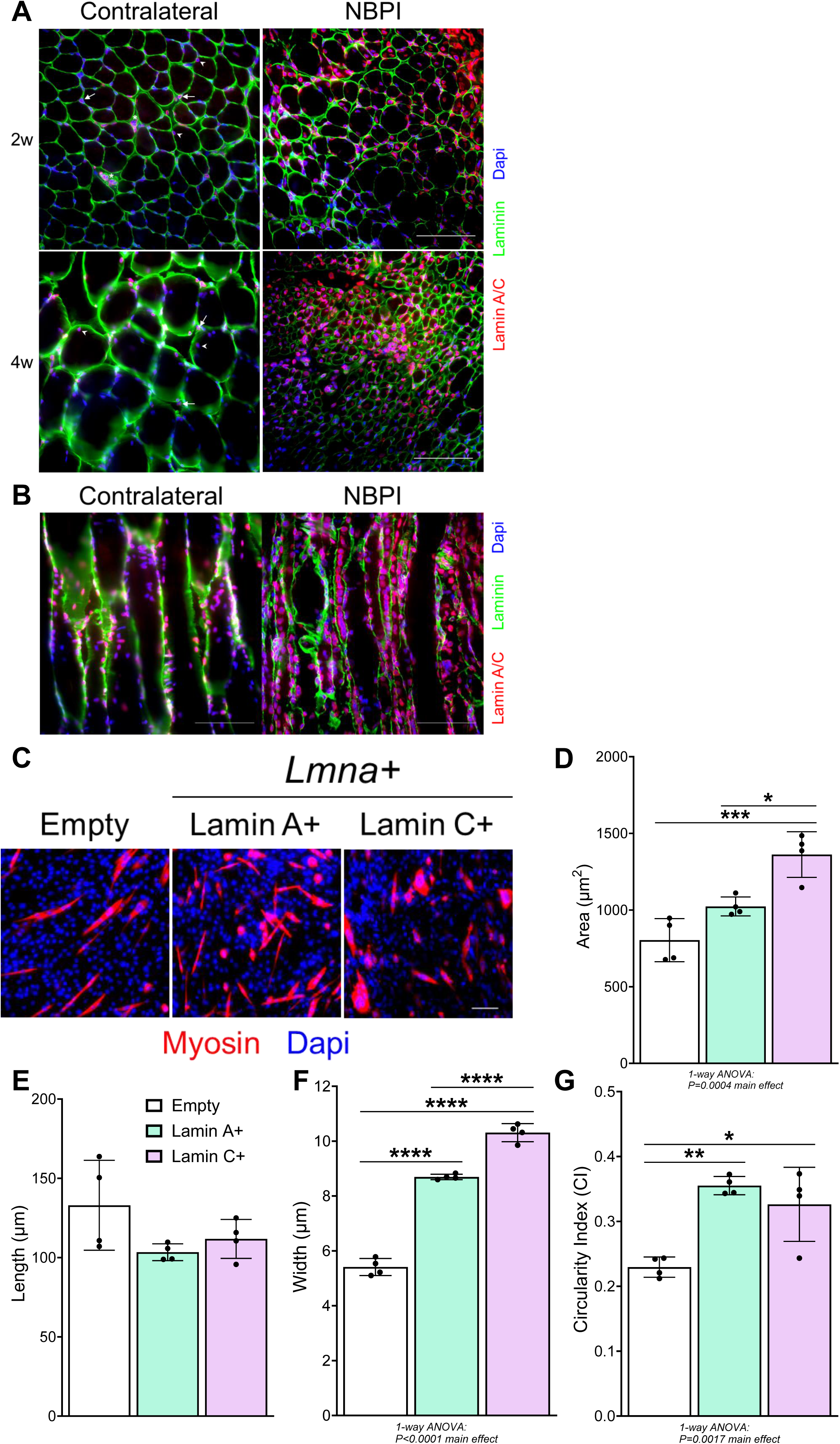
*Lmna* localization in vivo and overexpression in vitro. Representative immunostaining of biceps muscle sections in (**A**) transverse and (**B**) longitudinal view showed Lamin A/C expression. During neonatal muscle growth (left panels; contralateral), Lamin A/C+ is expressed by nuclei outside the myofiber (arrow), including nuclei in blood vessels (asterisk), and in peripherally located myonuclei within the myofiber (arrowhead). Following neonatal muscle denervation (right panels; NBPI), Lamin A/C is mainly expressed in centrally located myonuclei throughout the length of a myofiber. (**C**) Representative immunostaining of C2C12 skeletal muscle cells at three days of differentiation and quantitative analysis of cellular (**D**) area, (**E**) length, (**F**) width, and (**G**) circularity revealed dysmorphic changes in both Lamin A and Lamin C overexpressing conditions, characterized by altered width and circularity in the absence of changes to length compared to empty controls. Data are presented as mean + SD, n = 4 biological replicates per cell line. Statistical analyses: (**D**), (**E**), (**F**), (**G**) one-way ANOVA for cell line with a Bonferroni correction for multiple comparisons. *p<0.05, **p<0.01, ***p<0.001, ****p<0.0001. (**A**), (**B**), (**C**): 100 µm.

## Discussion

We previously showed that contractures are associated with sarcomere overstretch in denervated muscle,^8–11^ a longitudinal growth deficit mediated through MSTN signaling.^23^ However, the precise intracellular mechanism(s) by which MSTN regulates contractures remain unknown, as treatment with a decoy receptor (ACVR2B-Fc) rescues muscle growth and contractures independent of alterations to canonical MSTN signaling, namely Smad 2/3 activity, Akt/mTOR phosphorylation, and MuRF-1/Atrogin-1 expression.^23^ Our present study investigated the potential contributions of ERK and JNK signaling, two non-canonical pathways downstream of MSTN signaling, in contracture formation after neonatal muscle denervation. Our findings revealed that only JNK activity is elevated after NBPI, and pharmacologic blockade of JNK activation with SP600125 rescues NBPI-induced contractures, delineating a regulatory role for JNK in contracture modulation. This discovery supports contemporary reports of JNK signaling in mediating assorted muscle disorders and secondary muscle pathologies, including spinal muscle atrophy (SMA) and cachexia. Indeed, pharmacologic JNK inhibition is observed to reduce myofiber atrophy and maintain motor performance in a mouse model of SMA,^41^ as well as attenuate muscle wasting and improve long-term muscle function in cachectic mice.^42^ Importantly, the efficacy of SP600125 in preventing NBPI-induced contractures adds the JNK cascade to a growing list of therapeutic targets for pharmacologic contracture prevention, which currently includes proteasome activity^18–20^ and MSTN signaling.^23^ Furthermore, while associated pharmacologic agents (proteosome inhibitors, ACVR2B-Fc, SP600125) have not been established as safe for clinical use in children with NBPI-induced contractures, they provide an array of molecular tools to mechanistically dissect contracture pathogenesis and develop novel treatment strategies.

The JNK cascade elicits multiple downstream effectors, including *DNMT1*-mediated genome-wide methylation.^34^ While we observed elevated *Dmnt1* levels in NBPI consistent with increased global DNA methylation in muscle contractures in cerebral palsy,^43^ they are not alleviated by JNK inhibition. Instead, our findings identify the AP-1 target gene, *Lmna*, as a potential novel mechanism for contracture pathophysiology. A potential role for *Lmna* in contractures after neonatal muscle denervation is supported by contracture phenotypes present in *LMNA*-related disorders, known as laminopathies.^44,45^ The *LMNA* gene encodes the nuclear envelope proteins, Lamin A and Lamin C, which play multiple roles in nuclear shape and rigidity, mechanosensing, signaling, and gene regulation.^46^ Human mutations in *LMNA*, yielding either dysfunctional or excess Lamin A/C proteins, can result in contracture development.^47^ Indeed, dysfunctional Lamin A/C underlies Emery-Dreifuss muscular dystrophy (EDMD),^48^ which is characterized by progressive muscle contractures beginning in early childhood,^49^ whereas toxic accumulation of Lamin A/C mutant proteins underlies Hutchinson-Gilford progeria syndrome,^50^ a premature aging disorder also characterized by progressive muscle contractures.^51^ However, despite the prominence of contractures in the laminopathies, little is known regarding their pathophysiology.

Similarly, the role of *Lmna* in NBPI induced contractures remains to be elucidated. In our surgical NBPI model, we observed the localization of Lamin A/C primarily to central nuclei inside the myofiber in denervated muscles. These Lamin A/C+ centrally located myonuclei are also unevenly spaced longitudinally, often stacked in close proximity to one another along the myofiber. Since misaligned/displaced myonuclei are hallmarks of muscle disorder and dysfunction,^52–54^ these defects in nuclear position may contribute to contractures in neonatally denervated muscles. Since we also observed that Lamin A/C overexpression during myogenesis distorts muscle cell shape, a potential consequence of *Lmna* overexpression in denervated muscles involves cellular dysmorphia. However, it should be noted that the mechanism of JNK signaling and *Lmna* overexpression is not limited to impairments of muscle elongation. In fact, pharmacologic JNK inhibition does not correct sarcomere overstretch, potentially explaining why the rescue in contractures with SP600125 is moderate compared to other pharmacologic agents we have previously utilized, especially bortezomib.^18,19^ Nonetheless, we still found a correlation between sarcomere overstretch and contracture severity that was in fact strengthened by JNK inhibition, suggesting that JNK inhibition reduces the contribution of other determinants of contracture severity, which remain to be elucidated. Additional work with genetic models is needed to further interrogate JNK signaling and *Lmna* function in contractures.

While pharmacologic JNK inhibition does not correct impairments in longitudinal muscle growth in denervated muscles, it blunts volumetric growth and overall mass of normally innervated neonatal muscles. At present, limited data exist regarding the role of MSTN/JNK signaling in postnatal muscle growth. In postnatal muscles, JNK knockout impairs the hypertrophic response to mechanical overload, whereas JNK activation suppresses canonical SMAD-mediated MSTN signaling and enhances muscle hypertrophy after resistance exercise.^55^ Our current data extend these key findings by highlighting the requirement of JNK signaling for muscle growth specifically during the neonatal period, a critical and narrow window commonly overlooked in development of skeletal muscle.^23,56^ Importantly, these collective advances reveal that JNK signaling is essential for muscle growth under normal conditions, which contrasts its pathological role during muscle disease and injury.^41,42^ Moreover, findings here allow us to add the JNK signaling cascade to an growing list of cellular mechanisms with putative regulatory roles in neonatal muscle growth, including NRG/ErbB signaling^57^ and β-adrenergic signaling.^17^

The following limitations are acknowledged in this current study. We did not account for potential sex differences, despite our prior findings and recognition of sex-specificity in various domains of skeletal muscle.^17,23,56^ It is also entirely possible that JNK activation in contractures are not MSTN-induced. We do not believe this is likely since it would posit a divergence between MSTN-dependent and JNK-dependent contractures. Further, we cannot discount other pathways and target genes besides *Lmna* downstream of JNK signaling, including those with undefined functions in skeletal muscle. Future studies must thoroughly investigate such candidates.

## Conclusion

Our current study indicates NBPI-induced contractures occur at least in part through the JNK signaling cascade and potentially its downstream target *Lmna*, revealing a novel intracellular mechanism for neuromuscular contracture pathophysiology. In addition, findings here provide several mechanistic insights into aberrant muscle growth and contracture pathogenesis. First, the efficacy of SP600125 in preventing NBPI-induced contractures adds JNK inhibition to a growing list of therapeutic targets for pharmacologic contracture prevention, which includes proteasome activity and MSTN signaling. Second, the co-localization of Lamin A/C mainly to central nuclei inside the myofiber of denervated muscles posits a critical role of *Lmna* in governing myonuclear position during conditions of muscle wasting, while cellular dysmorphia observed with Lamin A/C overexpression during myogenesis posits a deleterious role for excess *Lmna* in muscle morphology. Additional *in vivo* and *in vitro* work are needed to further interrogate *Lmna* function in contractures and muscle growth. Third, the rescue of contractures in the absence of sarcomere length restoration with SP600125 indicates longitudinal muscle growth is not the sole determinant of contracture pathology. We must therefore account for additional factors, including muscle cell shape, and previously identified factors in future investigations to develop effective contracture therapies. Lastly, the requirement of JNK signaling for neonatal muscle growth provides greater mechanistic insights into muscle mass regulation, and potentially leads to a novel target for muscle restorative strategies.

## Materials and Methods

### NBPI surgical mouse model

Unilateral NBPI was surgically created in postnatal day (P) 5 wildtype mice (Charles River) by unilateral extraforaminal excision of the C5–T1 nerve roots under isoflurane anesthesia.^17,20,23^ This results in complete denervation of the forelimb and reliably causes elbow flexion contractures,^8–10^ which serves as our model for contracture investigation. Following surgery, mice were returned to their mums and housed in standard cages with bio-huts on a 1:1 light/dark cycle, with nutrition and activity *ad libitum*. Deficits in motor function were evaluated postoperatively and prior to sacrifice to ensure permanent deficits and prevent any confounding consequences of reinnervation. Mice displaying preserved or recovered movement in the denervated forelimb were subsequently excluded. Based on these criteria, we excluded 1 mouse from the study.

### Pharmacologic inhibition of JNK signaling pathway

In neonatal mice, JNK activity was inhibited with SP600125^36,37^ at 15mg/kg (Selleckchem #S1460). Drug treatments were administered via intraperitoneal injections beginning immediately after NBPI surgery at P5 and then every other day until harvest at two or four weeks post-operatively. Separate litters of mice were injected with 1% Dimethyl sulfoxide (DMSO) in phosphate-buffered saline (PBS) as controls. For four-week treatments, this regimen was performed twice to obtain two separate litters in both conditions. All control and experimental groups in this study were randomized by litter, with all treatments administered at noon and in the respective cages. To ensure adequate nursing and monitor for signs of toxicity, non-weaned mice were checked daily for milk spots, and daily body weight at all ages were recorded. Mice displaying an inability to nurse/self-feed post weaning were euthanized immediately.

### Assessment of contractures

At two weeks (P19) or four weeks post-NBPI (P33), mice were euthanized by exposure to overdose of inhalant anesthetic, 300µl Isoflurane in a drop box, until death is determined by cessation of respiration and heartbeat. A physical secondary method, either cervical dislocation (P19) or exsanguination (P33), was subsequently performed to ensure irreversible death. Following euthanasia at four weeks post-NBPI, bilateral elbow passive range of motion was assessed through a validated digital photography technique to determine elbow contractures, as previously described.^17,20,23^ Briefly, images of bilateral elbows were captured at maximum passive extension, and elbow flexion contracture severity was subsequently calculated in Zen (Zeiss) as the difference in passive elbow extension, paired between denervated and contralateral limbs. All digital photography and image analyses were performed with blinding to treatment groups. While representative images of forelimbs shown in **Figure 2B** have been processed to reflect comparable levels of sharpness, brightness, and contrast for illustrative purposes, no image manipulation was performed prior to measurements.

### Muscle tissue collection and processing

Following euthanasia at two weeks post-NBPI, and following digital photography for contracture measurement at four weeks post-NBPI, bilateral biceps muscles were harvested, flash frozen, and stored at –80°C for molecular analysis of ERK and JNK signaling, as well as downstream effectors of the JNK cascade. In separate litters of mice at two and four weeks post-NBPI, bilateral biceps muscles were embedded in optimal cutting temperature (OCT) compound, frozen in liquid nitrogen-cooled 2-methylbutane, cut at 10 µm for both transverse and longitudinal sections, and processed for immunofluorescence of Lamin A/C expression. At four weeks post-NBPI, the remaining forelimbs were positioned on cork at 90° elbow flexion, imaged by digital radiographs for humerus length, and fixed in 10% formalin for 48 h. Bilateral brachialis muscles were subsequently dissected, weighed, soaked in 25% Lugol solution (Sigma-Aldrich #32922) overnight, and processed for micro-computed tomography (Micro-CT) for whole muscle size on a Siemens Inveon PET/SPECT/CT scanner (Siemens Medical Solutions), as previously described.^17,23^ Post scanning, whole muscles were recovered in PBS overnight at 4°C and subsequently digested in 15% sulfuric acid for 30 min, whereupon muscle bundles were isolated for imaging of sarcomeres via differential interference contrast (DIC) microscopy at 40x on a Nikon Ti-E SpectraX widefield microscope.^17,20,23^

### Muscle morphometrics

Muscles with fewer sarcomeres in series require each sarcomere to overstretch to accommodate a given position, indicating a functional deficit in muscle length. We thus assessed sarcomere overstretch at fixed joint angles as an assay for serial sarcomere number or functional muscle length. Following microscopy acquisition of six different muscle bundles from the same muscle, brachialis sarcomere length was determined by measuring a series of 10 sarcomeres from each of the six representative DIC images in Zen.^17,20,23^ Humerus lengths were determined from digital radiographs by measuring the distance between the proximal humerus physis to the distal articular surface in Zen.^17,23^ Brachialis whole muscle measurements were obtained by processing the MicroCT scans into digital imaging and communications in medicine (DICOM) images with Fiji programs, specifically with Segmentation Editor to quantify whole muscle cross-sectional area (CSA), and with 3D Viewer to calculate muscle volume.^17,23^ Both CSA and volume measurements were subsequently normalized to humerus lengths of corresponding limbs. All quantitative measurements were performed with blinding to treatment. For illustrative purposes, representative sarcomere images shown in **Figure 3A** have been cropped to identical sizes, and processed to reflect comparable levels of sharpness, brightness, and contrast; whereas raw DICOM files were processed in Imaris software (Bitplane, Zurich, Switzerland) to create cross-sectional and whole-muscle images in **Figures 4A**. No image manipulation was performed prior to measurements.

### Muscle cell cultures

To examine the role of Lamin A/C during myogenesis, skeletal muscle cells (C2C12; ATCC) were plated in 6-well plates, and grown in Dulbecco’s Modified Eagle Medium (DMEM; Gibco) containing 10% fetal bovine serum (Hyclone^TM^). Proliferating C2C12 cells were transduced with a lentivirus coding for doxycycline-inducible Lamin A (Lamin A+) or Lamin C (Lamin C+) isoform of *LMNA*,^58^ or an empty cassette (Empty), with 6 µg/ml polybrene (Sigma-Aldrich) and subjected to spinfection by centrifugation at 600 x g for 1 h at 22°C immediately following viral overlay^59^ (all viral constructs were donated by Dr. Kohta Ikegami from Cincinnati Children’s Hospital Medical Center). All cell lines were incubated overnight and selected with antibiotic blasticidin (2 µg/ml; InvivoGen) for 48h prior to differentiation experiments. To induce differentiation, cells were first plated in 12-well plates, and switched to media containing DMEM and 2% horse serum (Hyclone^TM^) upon reaching 90% confluence. Doxycycline (1 µg/ml; Sigma-Aldrich) was added one day (24h) post-differentiation to induce gene expression. After three days (72h) of differentiation, all cell lines were processed for immunofluorescence, and parameters of cell shape were quantitated in Imaris (Bitplane).

To assay for Lamin A/C expression in vitro, cells were scraped in cold PBS, pelleted at 2,000 rpm for five minutes, and stored at –80°C for subsequent western blot. Non-transduced C2C12 cells were collected during proliferation and throughout day one (24h) till day six (144h) of differentiation, whereas transduced cells were collected at 36h of differentiation. A separate culture of non-transduced, proliferating C2C12 cells was treated with 50 µM Cisplatin (Enzo; #ALX-400-040) upon reaching 90% confluency, and collected after 16h of treatment as a positive control for cleaved Caspase-3.^60^

### Histological analyses

Cryosections were fixed in 1% PFA/PBS, permeabilized with 0.2% Triton X-100/PBS, blocked with 1% BSA/PBS/1% heat-inactivated donkey serum/0.025% Tween20/PBS, and incubated overnight at 4°C with anti-Lamin A (1:250; Abcam #ab26300) and anti-laminin (1:250; Abcam #ab14055). Following 1h of incubation with secondary Alexa Fluor antibodies (1:400) (Jackson ImmnunoResearch), slides were mounted with VectaShield containing DAPI (Vector Laboratories). Cell cultures were fixed in 70% methanol/30% acetone, permeabilized with 0.2% Triton X-100/PBS, blocked with 3%BSA/PBS, incubated overnight at 4°C with anti-sarcomeric myosin heavy chain (MHC) (1:20; clone MF20; Developmental Studies Hybridoma Bank), incubated for 1h with a secondary Alexa Fluor antibody (1:200) (Invitrogen), and rinsed in 0.1% Hoechst (Invitrogen)/PBS. All samples were visualized with a Nikon Ti-E SpectraX widefield microscope. For in vivo assessment of Lamin A/C expression, both transverse and longitudinal sections were visualized at 40X. For illustrative purposes, representative biceps muscle sections shown in **Figures 6A** and **6B** have been cropped to identical sizes, and processed to reflect comparable levels of sharpness, brightness, and contrast. To assay for muscle cell shape, five fields per well were first visualized at 10X with a standardized approach to image capture. The first image was taken at the center of the well, while the second, third, fourth, and fifth images were taken two fields of view (400 μm), respectively, to the right, top, left, and bottom of the first image, collectively depicting a square in the center of the well. The different parameters of muscle cell shape, including width, length, area, and circularity, were subsequently quantified through a binary thresholding algorithm developed in Imaris (Bitplane), with all analyses performed blinded to treatment. While representative muscle cell cultures shown in **Figure 6C** have been cropped to identical sizes, and processed to reflect comparable levels of sharpness, brightness, and contrast, no image manipulation was performed prior to measurements.

### Gene ontology

To search for relevant downstream JNK signaling targets, we began by identifying putative AP-1 transcriptional targets using the MotifMap transcription factor binding motif analysis.^38^ We then filtered these AP-1 target genes by human muscle phenotypes using the Enrichr gene set search tool,^39,40^ resulting in the following six genes: *ATL1*, *FAM126*, *FGD4*, *FLNC*, *KIF1A*, and *LMNA*. We proceeded to assess mRNA and/or protein expression of these candidate genes following NBPI and SP600125 treatment.

### Western blot

To explore signaling pathways downstream of MSTN in contracture formation and assess the effects of JNK inhibition on putative targets, we assayed for ERK and JNK signaling, as well as Caspase-3 and Lamin A/C expression, respectively, by western blotting as described previously.^23,61^ Briefly, bilateral biceps were bead homogenized in tissue lysis buffer [10 mM Tris (pH 7.4), 1 mM ethylenediaminetetraacetic acid (EDTA), 1 mM dithiothreitol, 0.5% Triton X-100, 2.1 mg/ml NaF] containing protease and phosphatase inhibitor cocktails (5 μl/ml; Sigma-Aldrich #P8340 and #P5726, respectively), whereas cell pellets were resuspended and sonicated in Chemicon lysis buffer [50 mM Tris (pH 6.8), 1 mM EDTA, 2% sodium dodecyl sulfate (SDS)]. Following centrifugation, the amount of protein in supernatants in muscle homogenates and cell lysates was quantified using Bradford protein assay, or BCA assay (Pierce^TM^), respectively. Samples were heated at 65°C for 30 minutes, separated on 10-12% SDS-PAGE gels (20 µg of proteins), and transferred at 4°C to PVDF-FL membranes (Immobilon-FL). Membranes were subsequently blocked in 5% dry milk or 5% BSA in Tris-buffered saline (TBS)-Tween, and incubated overnight at 4°C with antibodies against phosphorylated ERK1/2 (Thr202/Tyr204) (1:1000; Cell Signaling #9101), total ERK1/2 (1:1000; Cell Signaling #9102), phosphorylated SAPK/JNK (Thr183/Tyr185) (1:1000; Cell Signaling #9251), total SAPK/JNK (1:1000; Cell Signaling #9252), cleaved Caspase-3 (1:1000; Cell Signaling #9661) and total Caspase-3 (1:1000; Cell Signaling #9662), or an antibody against full length Lamin A and Lamin C (1:1000; Cell Signaling #2032). GAPDH (1:5000; Cell Signaling #2118) served as a control for sample loading. Membranes were then washed and incubated with a DyLight® anti-rabbit secondary antibody (1:5000; Cell Signaling #5151). Western blot signals were subsequently imaged on an Odyssey® CLx fluorescence scanner (LI-COR Biosciences). The relative abundance of phosphorylated and total protein levels of ERK and JNK were quantified using the Image Studio Lite program (LI-COR Biosciences), and normalized to each other and to corresponding GAPDH protein levels, with blinding to treatment groups.^23,61^ Nonphosphorylated and phosphorylated cell extracts for p44/42 MAPK (Erk1/2) (Cell Signaling #9194) and SAPK/JNK (Cell Signaling #9253), as well as muscle homogenates from adult mouse overloaded plantaris^23,61^ were used as controls for ERK and JNK signaling.

### RNA analyses

To assess activation of target genes downstream of JNK signaling, we performed RNA analysis as previously described.^61,62^ Briefly, total RNA was extracted from biceps muscle with TRI Reagent® (Sigma-Aldrich), and cDNA was synthesized using MultiScribe^TM^ reverse transcriptase with random hexamer primers (Applied Biosystems). Gene expression was assessed using standard qPCR approaches with PowerUp^TM^ SYBR Green Master Mix (Applied Biosystems). Analysis was performed on a CFX96™ System (Bio-Rad) with primers synthesized by Integrated DNA Technologies for the following genes: *Atl1*; *Dnmt1*; *Fam126a*; *Fgd4*; *Flnc*; *Kif1a*; and *Lmna* (**Supplemental Table 1**).

### Statistical analysis

All statistical analyses were conducted in Prism 10 software (GraphPad). Outliers in all datasets were detected *a priori* by Grubb’s test and excluded from subsequent analyses. The normality of all non-excluded data was then assessed by Shapiro-Wilk test. Unpaired two-tailed Student’s t-tests or Mann-Whitney U tests were used to compare normally distributed and non-normally distributed data, respectively, between animals. Paired two-tailed Student’s t-tests or Wilcoxon signed rank tests were used to compare normally distributed and non-normally distributed data, respectively, between denervated and contralateral forelimbs in individual animals. For data sets with two independent variables (NBPI surgery and pharmacologic treatment), a 2-way analysis of variance (ANOVA) with repeated measures between forelimbs and Bonferroni correction for multiple comparisons was performed. Data presented in this study are shown as mean ± standard deviation in all figures, with the level of significance between datasets depicted as: *p<0.05,**p<0.01,***p<0.001,****p<0.0001. A total of 82 mice were used in this study.

### Ethical statement

This study strictly adhered to the National Institutes of Health’s Guide for the Care and Use of Laboratory Animals, Animal Research: Reporting of In Vivo Experiments (ARRIVE) 2.0 guidelines, and Cincinnati Children’s Hospital Medical Center’s approved institutional animal care and use committee (IACUC) protocols (#2023-1010). Every effort was made to minimize suffering.

### Material Availability Statement

All data generated or analyzed during this study are included in the manuscript and supporting files.

## Supporting information

ARRIVE Checklist 2.0

## Acknowledgements

We thank the following entities within Cincinnati Children’s Hospital Medical Center (CCHMC): the Veterinary Services Core for surgical assistance, the Confocal Imaging Core for microscope assistance, and Melanie Gucwa from the Ikegami Laboratory for assistance in preparation of the viral constructs. We also thank Sharon Wang from the Preclinical Imaging Core (University of Cincinnati College of Medicine) for MicroCT assistance. This work was supported by grants to RC from the National Institutes of Health (NIH) (R01HD098280-01), as well as funding from the Division of Orthopaedic Surgery at CCHMC. DB was supported by the Child and Adolescent Health Medical Student Scholars Program through the Division of Pulmonary Medicine at CCHMC. The respective funding sources were not involved in the study design; in the collection, analysis, and interpretation of data; in the writing of the report; and in the decision to submit the paper for publication.

## Author Contributions

QG and RC conceived and supervised the study; KS, MG, DB, SC, QG, and RC designed the research and experiments; KS, MG, DB, SC, AT, KS-W, QG, and RC performed experiments; KS, MG, DB, SC, AT, GV and KS-W curated data; KS, MG, DB, SC, AT, GV, KS-W, QG, and RC analyzed data; QG wrote the manuscript with assistance from all authors.

**Table 1.** Key resources table.

| REAGENT or RESOURCE | SOURCE | IDENTIFIER |
| --- | --- | --- |
| **Antibodies** |  |  |
| Rabbit polyclonal Phospho-p44/42 MAPK (Erk1/2) (Thr202/Tyr204) | Cell Signaling Technology | Cat#9101 |
| Rabbit polyclonal p44/42 MAPK (Erk1/2) | Cell Signaling Technology | Cat#9102 |
| Rabbit polyclonal Phospho-SAPK/JNK (Thr183/Tyr185) | Cell Signaling Technology | Cat#9251 |
| Rabbit polyclonal SAPK/JNK | Cell Signaling Technology | Cat#9251 |
| Rabbit polyclonal Cleaved Caspase-3 (Asp175) | Cell Signaling Technology | Cat#9661 |
| Rabbit polyclonal Caspase-3 | Cell Signaling Technology | Cat#9662 |
| Rabbit polyclonal Lamin A/C | Cell Signaling Technology | Cat#2032 |
| Rabbit monoclonal GAPDH | Cell Signaling Technology | Cat#2118 |
| Goat anti-rabbit IgG (H+L) (DyLight® 800 4X PEG Conjugate) | Cell Signaling Technology | Cat#5151 |
| Chicken polyclonal anti-Laminin | Abcam | Cat#ab14055 |
| Rabbit polyclonal anti-Lamin A | Abcam | Cat#ab26300 |
| Alexa 488 donkey anti-chicken | Jackson ImmunoResearch | Cat#703-546-155 |
| Alexa 594 donkey anti-rabbit | Jackson ImmunoResearch | Cat#711-585-152 |
| Mouse monoclonal anti-MyHC | DSHB | Cat#MF-20 |
| Alexa 555 donkey anti-mouse IgG | Invitrogen | Cat#A-31570 |
| **Chemicals, peptides, and recombinant proteins** |  |  |
| SP600125 (Nsc75890) JNK inhibitor | Selleckchem | Cat#S1460 |
| Isoflurane, USP | Covetrus | NDC:1169567772 |
| Lugol solution | Sigma-Aldrich | Cat#32922 |
| DMEM, high glucose | Gibco | Cat#11965092 |
| Fetal bovine serum | Sigma-Aldrich | Cat#12306C |
| Horse serum | Gibco | Cat#16050130 |
| Penicillin/Streptomycin | Gibco | Cat#15070063 |
| Polybrene | Sigma-Aldrich | Cat#TR-1003-G |
| Blasticidin | InvivoGen | Cat#ant-bl-05 |
| Doxycycline Hydrochloride, ready-made solution | Sigma-Aldrich | Cat#D3072 |
| Cisplatin | Enzo Biochem | Cat#ALX-400-040 |
| Donkey serum | Jackson ImmunoResearch | Cat#017-000-121 |
| VectaShield with DAPI | Vector Laboratories | Cat#H-1200-10 |
| Hoechst 33342 | Invitrogen | Cat#H3570 |
| Protease inhibitor cocktail | Sigma-Aldrich | Cat#P8340 |
| Phosphatase inhibitor | Sigma-Aldrich | Cat#P5726 |
| Protein assay dye | Bio-Rad | Cat#5000006 |
| PVDF-FL membrane | Sigma-Aldrich | Cat#IPFL10100 |
| p44/42 MAPK (Erk1/2) Control Cell Extracts | Cell Signaling Technology | Cat#9194 |
| SAPK/JNK Control Cell Extracts | Cell Signaling Technology | Cat#9253 |
| TRI Reagent® | Sigma-Aldrich | Cat#T9424 |
| **Critical commercial assays** |  |  |
| Pierce™ BCA protein assay kit | Thermo Scientific | Cat#P122622 |
| High-Capacity cDNA Reverse Transcription kit | Applied Biosystems | Cat#4368814 |
| PowerUp™ SYBR™ GreenMaster mix | Applied Biosystems | Cat#A25742 |
| **Experimental models: Cell lines** |  |  |
| Mouse: myoblast cells | ATCC | Cat#CRL-1772 |
| **Experimental models: Organisms/strains** |  |  |
| CD-1® IGS mouse | Charles River | Cat#022 |
| **Oligonucleotides** |  |  |
| See Supplemental Table 1 for all qPCR primers | Integrated DNA Technologies | N/A |
| **Recombinant DNA** |  |  |
| pCW57-MCS1-P2A-MCS2-PGK-Blast | This paper | Ikegami lab ID: bKI214 |
| Wild-type Lamin A cDNA in pCW57-MCS1-P2A-MCS2-PGK-Blast | Ikegami et al.<sup>58</sup> | Ikegami lab ID: bKI218 |
| Wild-type Lamin C cDNA in pCW57-MCS1-P2A-MCS2-PGK-Blast | Ikegami et al.<sup>58</sup> | Ikegami lab ID: bKI246 |
| **Software and algorithms** |  |  |
| Zen v3.3 (blue edition) | Zeiss | RRID:SCR_013672 |
| NIS-Elements | Nikon Instruments | RRID:SCR_014329 |
| Imaris software | Bitplane | RRID:SCR_007370 |
| Fiji-ImageJ | Schindelin et al.<sup>63</sup> | RRID:SCR_002285 |
| MotifMap | Xie et al.<sup>38</sup> | N/A |
| Enrichr | Chen et al.<sup>39</sup> | RRID:SCR_001575 |
| LI-COR Image Studio Acquisition software | LI-COR Biosciences | RRID:SCR_015795 |
| CFX Maestro software | Bio-Rad | RRID:SCR_018064 |
| GraphPad Prism v10 | GraphPad Prism Inc. | RRID:SCR_002798 |

## Abbreviations

NBPI: neonatal brachial plexus injury
MSTN: myostatin
ERK: extracellular receptor kinase
JNK: c-Jun N-Terminal kinase
*DNMT1*: DNA methyltransferase 1
*FHL1*: four and a half LIM domains 1
*FLNC*: filamin C
*LMNA*: lamin A/C
GAPDH: glyceraldehyde-3-phosphate dehydrogenase
DPBS: Dulbecco’s phosphate-buffered saline

---

**Figure 1-supplemental figure:**
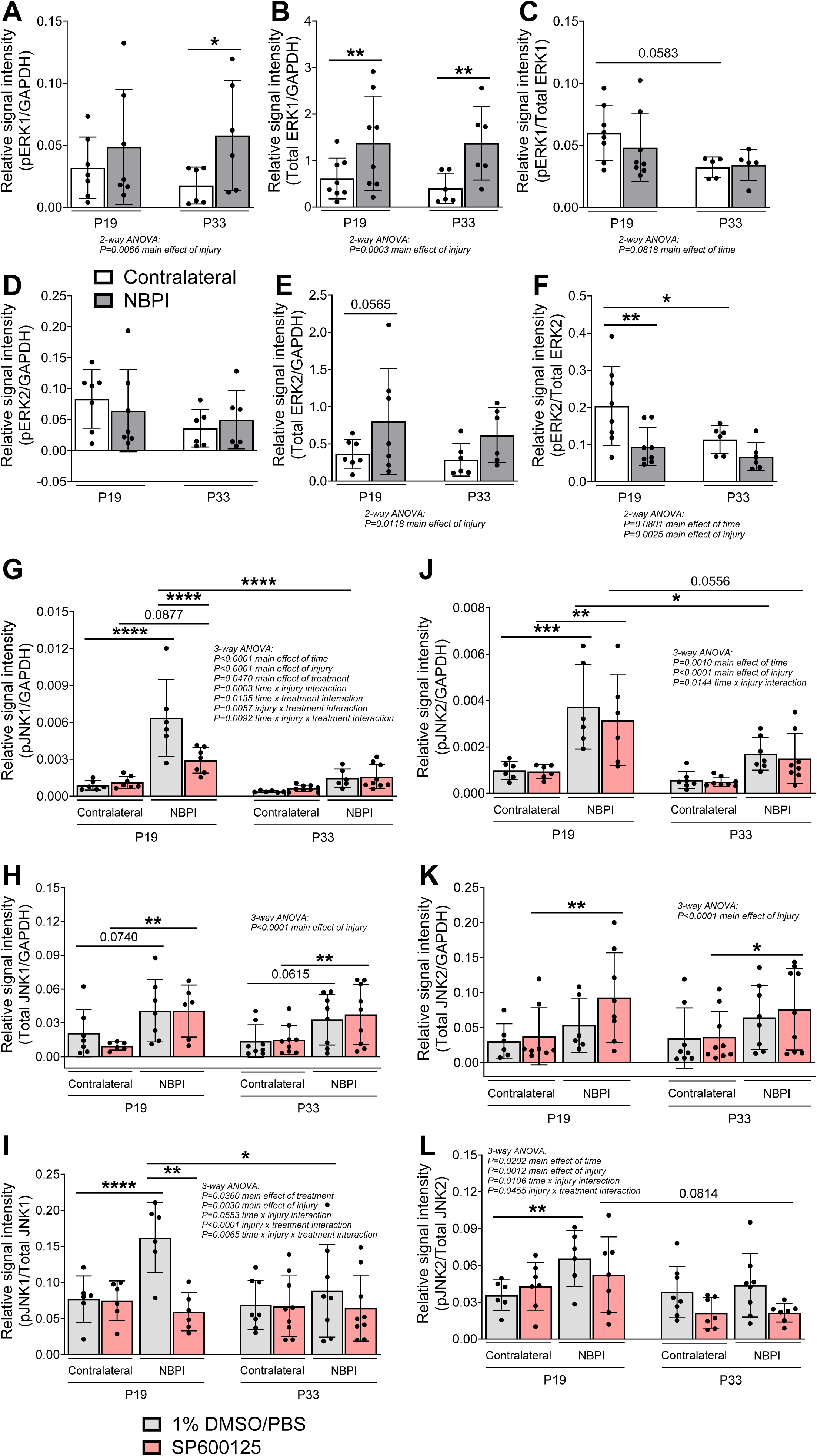
Isoforms of ERK and JNK pathways after NBPI. Neonatal denervation increases (**A**) ERK1 phosphorylation (pERK1) and (**B**) ERK1 translation in biceps brachii muscles. However, (**C**) the normalized pERK1 western signal, along with (**D**) phosphorylated ERK2 (pERK2), (**E**) total ERK2, and (**F**) the normalized pERK2 western signal, are not elevated by NBPI. Data are presented as mean ± SD, n = 6-8 independent mice per group. In contrast, at two weeks post-NBPI, neonatal muscle denervation upregulates both (**G**) phosphorylated JNK1 (pJNK1) and (**J**) phosphorylated JNK2 (pJNK2), as well as the (**I**) normalized pJNK1 and (**L**) pJNK2 western signals, without altering (**H**) JNK1 or (**K**) JNK2 translation. Moreover, pharmacologic inhibition of JNK with SP600125, specifically (**G**) blunts JNK1 activation and (**I**) restores JNK1 signaling to basal levels. Data are presented as mean ± SD, n = 6-9 independent mice per group. Statistical analyses: (**A**), (**B**), (**C**), (**D**), (**E**), (**F**) two-way ANOVA for NBPI surgery (repeated measures between forelimbs) across time with a Bonferroni correction for multiple comparisons, (**G**), (**H**), (**I**), (**J**), (**K**), (**L**) three-way analysis of variance (ANOVA) for pharmacologic treatment and NBPI surgery (repeated measures between forelimbs) across time with a Bonferroni correction for multiple comparisons. *p<0.05, **p<0.01, ***p<0.001, ****p<0.0001.

**Figure 4-supplemental figure:**
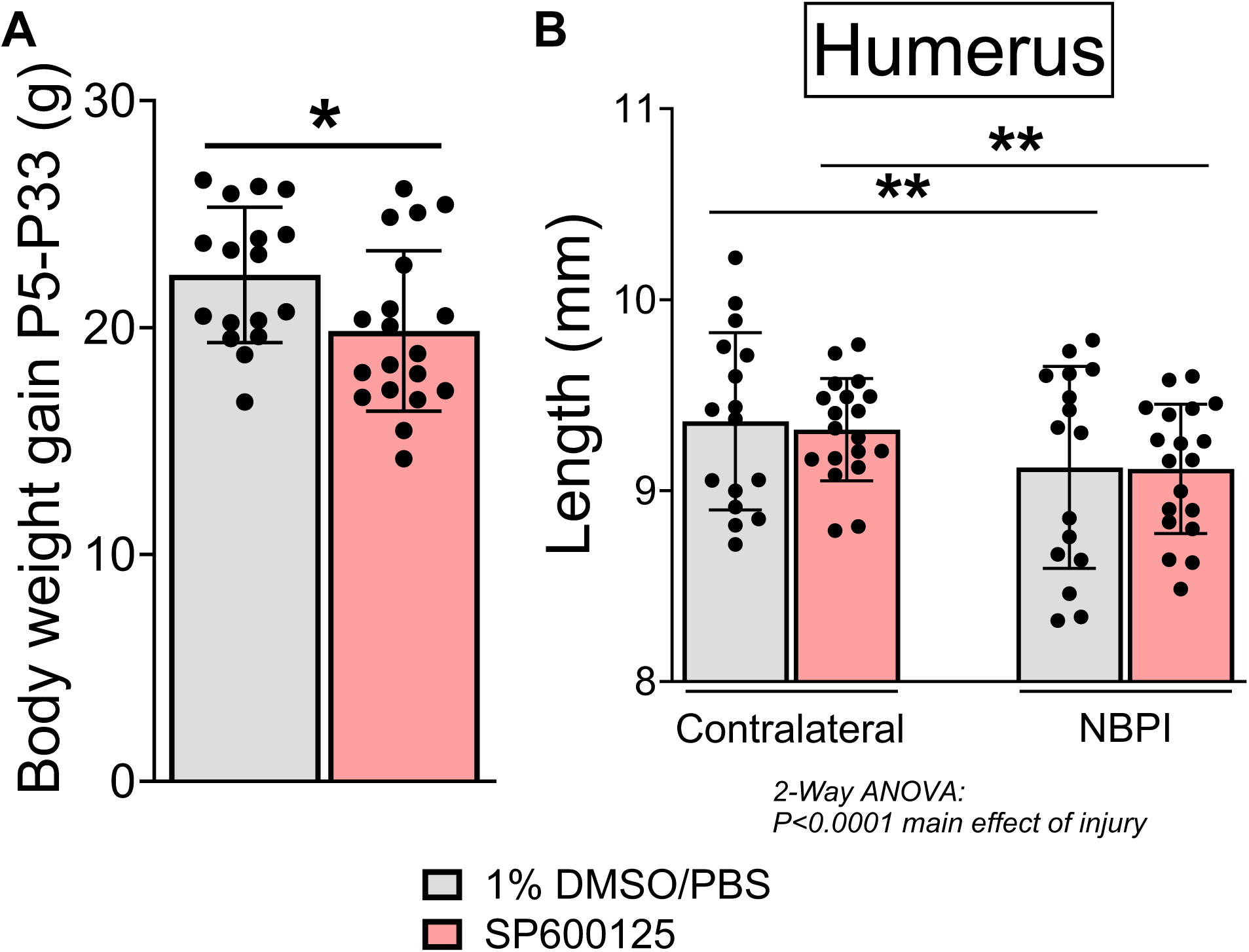
Off-target effects of pharmacologic JNK inhibition. SP600125 treatment (**A**) impeded additional weight gains during developmental growth, (**B**) but did not alter humerus lengths in either denervated (NBPI) or non-operated (contralateral) forelimbs. Data are represented as mean ± SD, n = 16-19 independent mice per group. Statistical analyses: (**A**) unpaired two-tailed Student’s t-tests between treatment groups, (**B**) two-way ANOVA for pharmacologic treatment and NBPI surgery (repeated measures between forelimbs) with a Bonferroni correction for multiple comparisons. *p<0.05, **p<0.01.

**Figure 6-supplemental figure:**
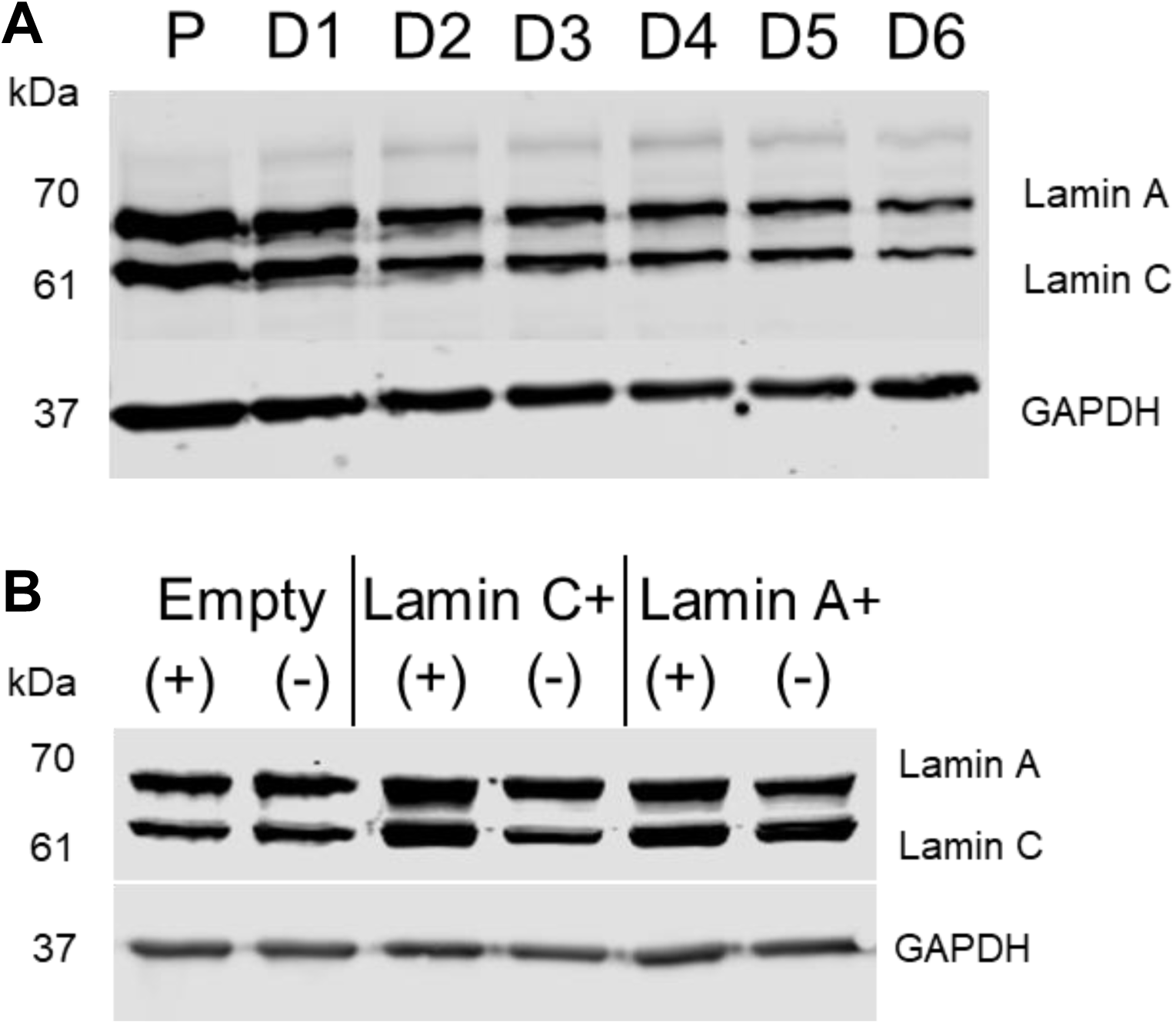
LMNA protein expression in vitro. LMNA protein levels in C2C12 skeletal muscle cells (**A**) at different stages of myogenesis during cellular proliferation (P) and throughout six days of cellular differentiation (D), and (**B**) at three days of differentiation in control (Empty), Lamin A (Lamin A*+*), or Lamin C (Lamin C*+*) overexpressing conditions, in the presence (+) or absence (-) of doxycycline.

**Supplemental Table 1.**
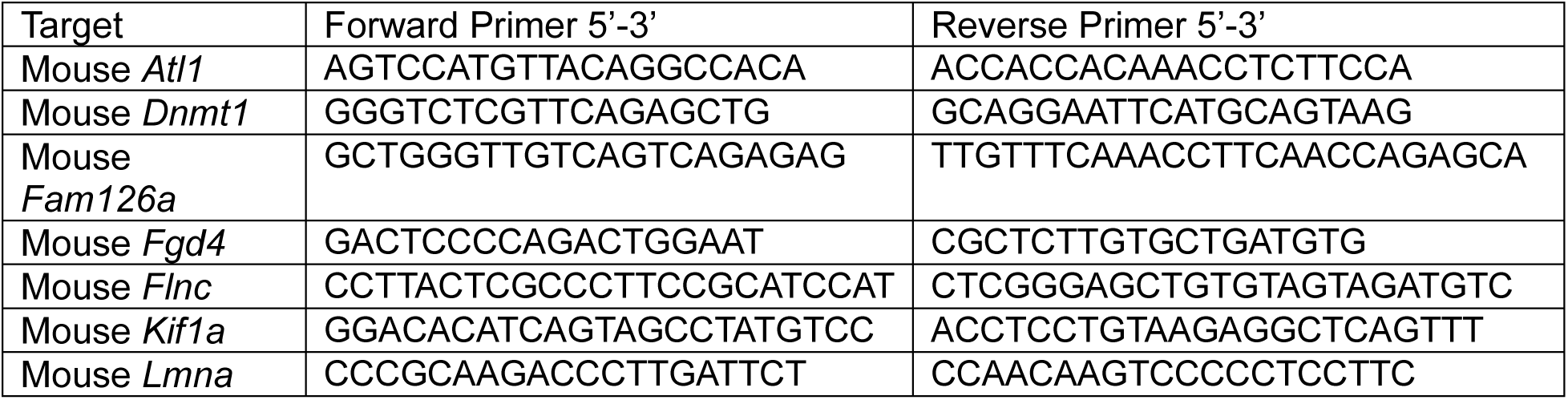
qPCR primers.

